# Comparison of Interventional Causal Structure Learning Algorithms for Gene Regulatory Network Inference

**DOI:** 10.64898/2025.12.05.692565

**Authors:** Jan L. Sprengel, Britta Velten

**Affiliations:** Centre for Organismal Studies, Heidelberg University, Im Neuenheimer Feld 230, Heidelberg, 69120, Germany; Interdisciplinary Center for Scientific Computing, Heidelberg University, Im Neuenheimer Feld 205, Heidelberg, 69120, Germany

**Keywords:** single cell CRISPR Screening, Gene Regulatory Networks, Causal Structure Learning

## Abstract

**Background:** Causal structure learning offers a promising approach to studying gene regulation in cells, aiming to provide deeper mechanistic insights than purely association-based methods. Theoretical groundwork indicating that interventions improve identifiability of causal structure motivates the use of causal structure learning methods in scenarios with interventional information.

**Results:** This benchmark investigates the ability of existing causal structure learning algorithms to leverage the information revealed by targeted interventions to infer gene regulatory networks (GRNs). In this study both synthetic and experimental single-cell CRISPR perturbation data is leveraged, and a suite of causal structure learning algorithms, is evaluated on metrics tailored to synthetic ground truth and real biological data respectively. On synthetic data, accurate recovery of GRNs is achieved under favourable conditions: strong interventions, large sample sizes, and low measurement noise. However, on real data, performance remains unreliable, limited by technical and biological noise, as well as algorithmic scalability. This highlights a current gap between theoretical potential and practical application of causal structure learning for GRNs.

**Conclusions:** The benchmark provides insight into algorithm strengths and limitations and offers groundwork for further methodological development. We also provide an accessible software package to leverage modern causal structure learning on custom datasets and thereby foster future exploration of the potential of causal structure learning.

## 1 Background

Understanding gene regulatory networks (GRNs) plays a central role in cellular biology and disease research, but a direct inference from observational data is challenging [11]. Advances in single-cell RNA sequencing (scRNA-seq) combined with CRISPR (clustered regularly interspaced short palindromic repeats) perturbations now make it possible to measure transcription responses to targeted gene interventions [3]. This opens the door to applying causal structure learning methods, originally developed in machine learning, to infer gene regulation from interventional data. While these methods show theoretical advantages, especially in improving the identifiability of causal structure [10, 20], they are often developed in the context of simplistic scenarios, and their ability to transfer to realistic biological settings remains unclear. In typical scCRISPR data, there are a number of challenges such as technical noise of the scRNA-seq process, inherent biological variability of gene expression, and possible cellular variation in perturbation efficiency [2, 12].

There exist previous benchmarks on GRN inference using observational gene expression data or multimodal data [11, 18]. For interventional data, CausalBench evaluated causal structure learning methods using both biological annotations and statistical metrics [6]. It highlights a poor performance of pre-existing theoretically founded causal models, which are observed to be outperformed by empirical approaches developed for the context of the benchmark.

In this work, we extend upon the work of CausalBench by adding a detailed evaluation of methods on synthetic data to better understand their performance gap when used with scRNA-seq data. We also test a number of different evaluation criteria on experimental data, and extend upon the selection of algorithms by using some of the highly-performing models of CausalBench, and adding previously unconsidered approaches such as a supervised model [13], and providing modifications of an existing algorithm to the distribution of RNA-seq data characteristics.

In addition, we provide scp-infer (single-cell perturbation inference), a well documented python package, which implements six causal structure learning methods including classical and neural approaches. scp-infer furthermore contains an extended version of the SERGIO framework [7] for the generation of synthetic gene expression data, as well as functionalities such as pre-processing and evaluation metrics for GRN inference with scCRISPR data. The toolkit thus significantly lowers the hurdle for researchers to apply interventional causal structure learning methods to scCRISPR data.

Using scp-infer, we benchmark six causal structure learning algorithms on synthetic data as well as real gene expression data from a multimodal Perturb-CITE-seq screen in patient-derived melanoma cells by Frangieh et al. [8] (Fig. 1). Our results provide a detailed comparative view of algorithms’ abilities to recover causal gene interactions under different practical constraints such as sample and network size, intervention availability, noise and measurement variability, and help to understand their strengths and limitations when applied to experimental data.

**Fig. 1.**
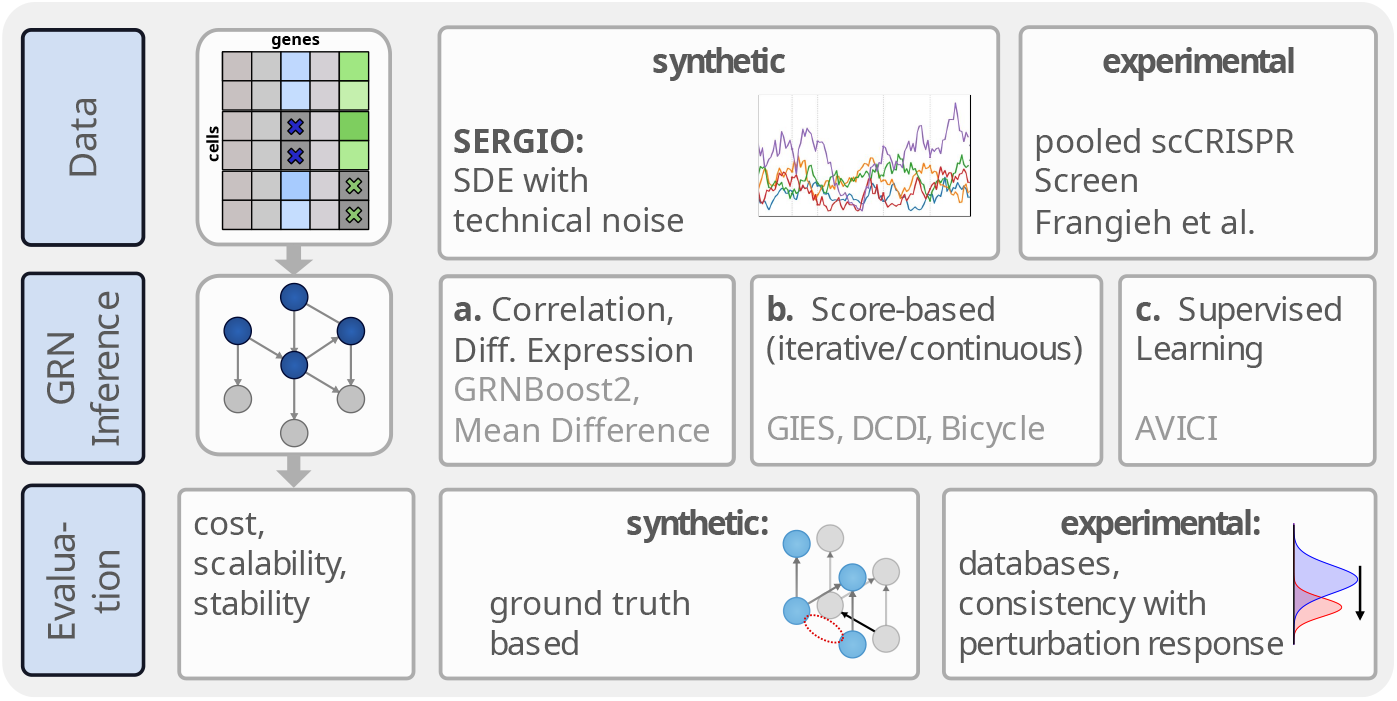
Overview of the type of data and methods for GRN inference and evaluation used in the Benchmark.

As new methods for interventional causal structure learning are developed scp-infer may easily be extended to include them.

## 2 Methods

### Data Generation

To generate synthetic gene expression data for the context of the benchmark we repurposed the SERGIO framework [7], integrating a Stochastic Differential Equation (SDE) for the concentration of each molecular species of RNA over time, and subsequently taking samples to produce a time-agnostic gene expression matrix 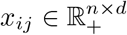, where *n* is the number of samples taken and *d* the number of genes considered. Knockdowns on selected genes were simulated by altering the SDE, and technical noise was modelled by transforming the observed samples *x*_*ij*_. (See Supplement E for details) In order to evaluate algorithms’ performance we selected a baseline scenario which for most algorithms proved to be manageable, and then modified individual parameters to observe how changes in data characteristics affect inference results. As a baseline scenario we simulated gene expression for 10 genes, which contain five underlying regulatory edges, randomly selected following an Erdős-Rényi model, and modelled knockouts on all 10 genes. From this baseline scenario we then altered parameters such as the number of samples *n*, the number of genes *d* or the number of distinct performed interventions. (Table E1, E2)

### Experimental Data

Alongside synthetic data, experimental RNA-seq data was used for model evaluation, for which we selected gene expression from a Perturb-CITE-seq screen of patient-derived melanoma cells on which 249 different genes were targeted with CRISPR-Cas9 perturbations [8]. The provided gene expression matrix was preprocessed with a standard sc-RNA-seq workflow; it was filtered for quality control, adjusted for the library size per cell and counts were transformed by the shifted logarithm [14]. We then selected a low number of 40 genes that were targeted in the screen for GRN inference, resulting in a matrix of dimensions *d* = 40 genes and *n* = 12064 cells (Table G3).

### Causal Model of Gene Regulation

In order to discuss the causal structure of GRNs, we will primarily make use of the framework of structural causal models (SCM) [17]. Here the genes are represented by a set of *d* variables **x** = (*x*_1_, …, *x*_*d*_), and their causal structure is represented by a directed graph *G*, whose edges correspond to all direct regulatory effects between the genes.

The generative process of **x** can then be denoted by a set of *d* general functions *f*_*i*_, each defining the value of *x*_*i*_ and only depending on the direct causes of *x*_*i*_, which correspond to its parents in *G*.

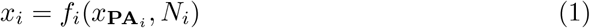

Where *f*_*i*_(..) is a general function that reflects the regulatory effects and *N*_*i*_ is an independent random variable which facilitates the introduction of stochastic effects. The parents 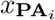 of *x*_*i*_ then correspond to all of the regulators of gene *i* in the GRN, while its children 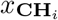 correspond to the regulatory targets. Within this framework targeted interventions on a gene *i* may be modelled by modifying the function *f*_*i*_ that defines the expression behaviour of the gene.

In general causal sufficiency is assumed, i.e. that **x** contains all common direct regulators of the considered variables *x*_*i*_. Some of the methods also assume that the GRN does not contain feedback loops, in which case the graph *G* of the SCM is constrained to a directed acyclic graph (DAG).

### Causal Structure Learning Algorithms and Application

In this benchmark we selected a number of algorithms, with a focus on algorithms that have a strong basis in causal modelling and are able to leverage interventional data, where we chose algorithms to represent different methodological approaches (Figure 1). As a classical algorithm we considered the score based iterative search algorithm GIES [10]. We considered two continuous optimisation methods DCDI [4] and Bicycle [19]. DCDI performs continuous optimisation of a causal model with an acyclic underlying graph and modifiable variable distributions, such as a Gaussian option. Bicycle is specifically designed for gene expression data, modelling gene expression as the steady-state of an Ornstein-Uhlenbeck process and allowing cyclic causal structures. Furthermore we applied the supervised deep-learning based algorithm AVICI [13]. As a top performer in the CausalBench benchmark the difference in mean expression, which we will refer to as ‘Mean Difference’ [5], and which simply considers average differential expression under interventions, was also applied. Lastly GRNBoost2, which is widely used in GRN inference, e.g. as part of the SCENIC framework, and does not explicitly leverage interventions, was used as a baseline [1, 15].

Of these algorithms Bicycle and AVICI contain design elements for scRNA-Seq data, and can be applied to the discrete read counts of each gene directly. GIES and DCDI however assume normally distributed data by default; we apply these to transformed gene expression data, where however the presence of zero-counts stands in conflict with their assumed bell-shaped distribution. As a possible augmentation to DCDI, we thus modified its likelihood function to extend it to non-Gaussian distributions characteristic to transcriptomics (See Supplement B for details). For one a Gaussian mixture model to operate on transformed reads was designed, where one Gaussian component of the mixture fixed at *µ* = 0 accounts for zero-counts. We experimented with ways to set the mixture rate, finally settling on a configuration where DCDI freely learns it as an additional variable of its likelihood function (DCDIDrop-local). Additionally DCDI based on a negative binomial distribution was applied directly on discrete reads (DCDI-NB).

For AVICI we applied two pre-trained models for which the weights are supplied by the authors. The first one, which was pre-trained with data from a variety of general SCM models (AVICI-raw), was applied on the synthetic data. The second one, which was trained on count synthetic gene expression data generated with SERGIO (AVICI-GRN), was applied on the synthetic data with technical noise and the experimental data.

When applying causal structure learning methods further considerations had to be taken into account per method. For the continuous optimisation models DCDI and Bicycle a range of hyperparameters with regards to training procedure and model architecture had to be specified — we here adopted the parameter choices recommended by the original authors where available and performed a grid search for remaining parameters to identify optimal values.

Unlike DCDI, GIES and AVICI, which predict the binary presence of edges, GRN-Boost2, Mean Difference and Bicycle produce a weighted interaction score between features. In order to gain comparable outputs across all methods, we therefore applied a threshold to the weighted interaction scores to obtain binary GRNs. Details on all considered methods can be found in Supplement A.

### Evaluation Metrics

In order to evaluate GRN predictions, a suite of metrics, tailored to synthetic and biological contexts respectively, were used. For synthetically generated gene expression data the ground truth (GT) GRN is known, which can then be compared against the predicted graphs to rate their quality - for this primarily the *F*_*β*_-Score, a weighted harmonic mean of precision and recall, is reported (Supplement D).

In the context of biological gene regulation no reliable GT is available. Here we used a proxy for the ground truth gene interaction in *homo sapiens*, the annotated GRN CollecTRI [16]. The Jaccard index was used to compare edge presence between this GRN and method predictions (See Supplement D).

Furthermore observed differential expression of genes under perturbations was leveraged to assay GRN quality. For one, the average differential expression of genes that are predicted children of a perturbed gene was computed, using the value of Wasserstein distance between their distribution in the perturbed and unperturbed context, as previously suggested by CausalBench [6] (See Supplement D).

Alternatively we report the portion of observed differentially expressed (DE) genes across perturbations, which can be explained by edges outgoing from the targeted gene in the predicted GRN. We use an adjusted t-test with a p-value threshold of 0.05 and multiple-testing correction to identify significant DE-genes under each perturbation, and compute the portion of these which is connected to the target of the perturbation in the predicted GRN considering different numbers of intermediate steps.

## 3 Results

### Synthetic Data

When observing the performance of the considered causal inference methods we notice a very strong dependence on simulation parameters and data characteristics.

In our baseline configuration, we observe high *F*_0.5_-scores with averages of 0.90 ± 0.10 for DCDI, 0.92 ± 0.09 for AVICI, 0.76 ± 0.13 for GIES and 0.80 ± 0.12 for Mean Difference. All these methods which can leverage interventional information outperform GRNBoost2, which often predicts edges with reversed orientation, producing a lower score of 0.47 *±* 0.05 (Fig. 2 a.,b.,c.).

**Fig. 2.**
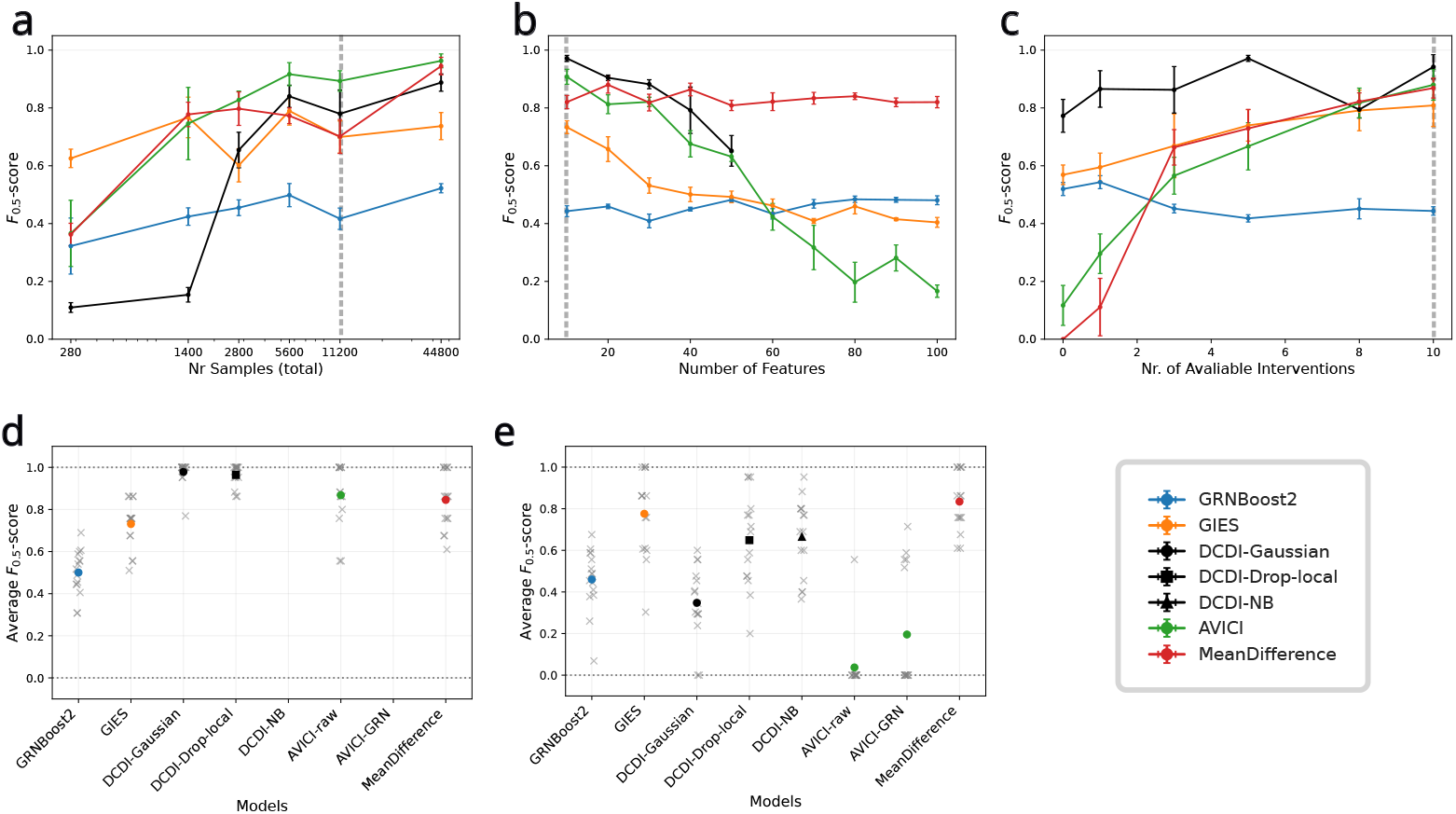
Synthetic Data: **a-c**. *F*_0.5_-Scores of the four algorithms on synthetic data without technical noise in different simulation scenarios; **a**. sample number, **b**. gene number and **c**. intervention number; values were computed over 5 separate datasets each, plotted is the mean and bars indicate standard error of the mean. The dashed vertical line marks the baseline configuration which we used for each parameter. **d and e**. *F*_0.5_-Scores of the algorithms on synthetic data without **d**. and with **e**. simulated scRNA-Seq technical noise, crosses are individual results on 15 separate datasets, coloured dots indicate the average. DCDI-NB and AVICI-GRN are not present in **d** as they require count data.

We observe that the prediction qualities largely improve with increasing samples, especially so for DCDI, which only shows competitive performance from 2,800 samples upwards (Fig. 1 a.). Likewise with an increasing number of available interventions the methods making use of interventions improve in performance (Fig. 1 c.) In contrast, with increasing number of genes the methods based on structural causal models struggle, producing either worse predictions, or in the case of DCDI not terminating for *d >* 50 within the time limit of 48h and 40 GB of available GPU Memory (Fig. 1b.). Only GRNBoost2 and Mean Difference do not loose performance from larger gene numbers. We conducted further tests, for which the results can be viewed in supplementary figure Fig. E6. We also varied the density of underlying GRNs, finding that GRNBoost2 struggles to identify the correct edge orientation and predicts many reverse GRN edges. In turn Mean Difference includes false positives (FP) where there are indirect regulation effects. DCDI and GIES also predict a number of incorrect edges while capturing a majority of the correct ones, whereas AVICI rarely predicts incorrect edges, but in some cases fails to capture existing ones (Fig. E7).

### Data with Technical Noise

When simulated technical noise is applied to the synthetic data we in turn observe a decrease in performance of some methods (Fig. 1 d. and e.). GRNBoost2, Mean Difference and GIES appear largely unaffected, while for DCDI and AVICI there are significant decreases in performance. DCDI with a Gaussian distribution achieves *F*_0.5_ values of 0.98 ± 0.05 without synthetic noise, but only 0.35 ± 0.18 with technical noise. The modified non-Gaussian versions maintain higher scores, of 0.65 ± 0.22 for DCDI-Drop-local and 0.66 ± 0.18 for DCDI-NB, see Supplement B Fig. B3 for other DCDI configurations. For AVICI both pretrained models tend to predict graphs that are overly sparse, typically featuring a precision comparable to other models but much lower recall, with some of the predicted GRNs being empty (see Fig. G9).

### Experimental Data

When transferring from synthetic to experimental data a marked shift in behaviour is visible. Now predicted GRNs are less consistent with very small overlaps; when computing overlaps between the predicted GRNs of different methods using the Jaccard Index we see overlaps below 0.35 between any of the predictions (Fig. 3 a).

**Fig. 3.**
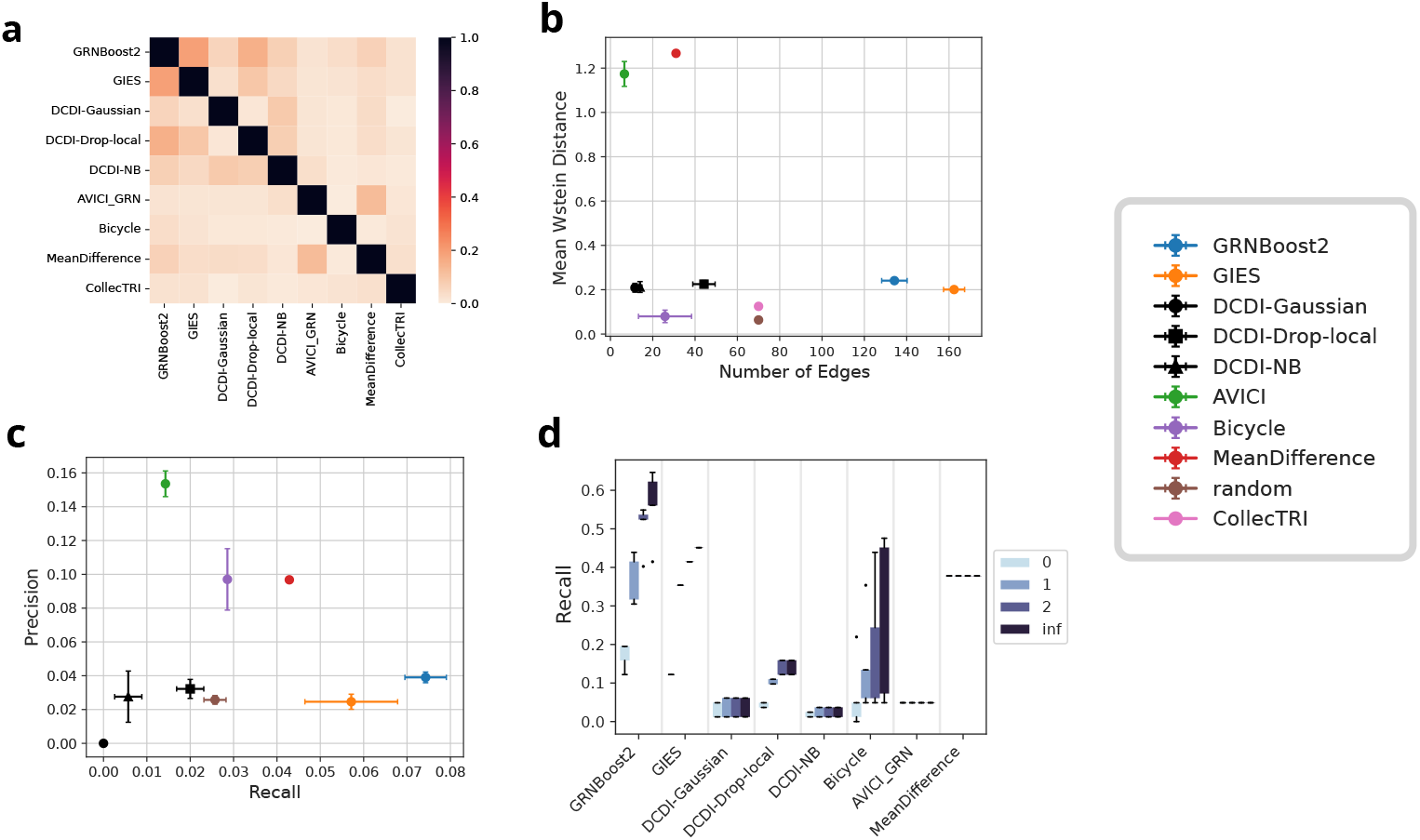
Experimental Data: **a**. Pairwise Jaccard indices between GRNs of each algorithm. **b**. Mean Wasserstein Distance plotted over the total number of edges in the GRN. **c**. Precision and Recall of GRNs with respect to CollecTRI grund truth. Markers indicate the mean and error bars the standard deviation over five independent runs. **d**. Recall of the inferred GRNs wrt. genes that showed differential expression under each perturbation, colour indicates the number of indirect steps considered, boxplot over the values of five independent runs.

Comparing the predictions across methods to statistical and biological baselines we observe an apparent precision-recall trade-off, with methods predicting drastically different numbers of GRN edges (Fig. 3 b,c). We observe that methods with low number of predicted edges, especially AVICI, tend to produce a high mean Wasserstein score (statistical measure) or precision (biological annotation) (Fig. 3 b,c). In a similar fashion e.g. GRNBoost2 predicts a large number of edges, but features a lower precision or mean Wasserstein score (Fig. 3 b). We note that for example Mean Difference features a comparatively high score in the statistical metric, but a lower score with respect to the biological annotation (Fig. 3 b,c), which is in agreement with the observations made by CausalBench in a different scCRISPR-Screen [6].

When analysing the recall of predicted GRN edges to gene pairs that show differential expression, we see a trend that methods which inferred dense GRNs, especially GRN-Boost2 and GIES, also achieve the highest recall values (Fig. 3 d). These are followed by Bicycle and Mean Difference, the latter of which achieves a recall almost as high as GIES despite containing only a portion of its edge numbers.

## 4 Discussion and Conclusions

Here we investigated the ability of methods rooted in structural causal discovery for inferring gene regulatory networks from interventional data to understand the extent to which their theoretical benefits could improve performance of state of the art methods for sc GRN inference.

Applying the methods in different contexts we found that interventional causal models faithfully reconstruct GRNs from synthetic data under good conditions, and can improve upon commonly used GRN inference approaches from observational single cell data. However these methods based on the structural causal model framework appear to be impeded significantly by a number of factors which are present in biological data. They are hampered by the biological and technical noise effects of RNA-sequencing, and struggle to scale to large network sizes, as a result of which the considered subsets do not necessarily obey the assumed causal sufficiency. In turn simpler empirical methods such as GRNBoost2 and the Mean Difference appear fairly robust to varying data features and technical noise, but as they do not feature the means for causal disentanglement, they do not achieve perfect reconstruction even in ideal conditions.

We recommend that biologists wishing to infer regulatory interactions from scRNA-seq technologies consider the presented GRN inference algorithms, but proceed with caution when interpreting results as no single currently available method is ideal for the task, and quantifying the confidence of predicted regulatory interactions remains difficult. The provided toolkit in python can help users to test various methods and evaluate them on custom datasets.

## Declarations

### Availability of data and materials

The processed experimental data used during this analysis is publicly available via the Single Cell Portal: https://singlecell.broadinstitute.org/single_cell/study/SCP1064 [9].

A utility to facilitate applying GRN inference Algorithms to custom scCRISPR-Screen data is available under https://github.com/velten-group/scp_infer, the full code to replicate the results of this study is available under https://github.com/velten-group/scp_infer_analysis.

### Authors’ contributions

The research was carried out by JS under supervision of BV.

## Acknowledgements

We would like to thank Purusharth Saxena for testing the software implementation and providing helpful feedback.

