## Appendix for "Comparison of Interventional Causal Structure Learning Algorithms for Gene Regulatory Network Inference"

### Part I
### Appendix

**B Adaptation of DCDI to Noise** **21**

**C Methods: Application of Causal Discovery Algorithms** **24**

**D Methods: Evaluation Metrics** **26**

**E Methods: Benchmark with Synthetic Data** **28**

**F Methods: Benchmark with Experimental Data** **33**
F.1 Data Preprocessing . . . . . . 33
F.2 Application of Causal Discovery Algorithms . . . . . . 35

**G Additional Results** **36**

### A Introduction of Causal Discovery Algorithms

We here present the general working principles of the considered causal structure learning methods. We include information on GENIE3 as this is what GRBoost2 is largely based on.

#### A.1 GENIE3

GENIE3 [15] is a tree-based tool for unsupervised GRN inference using gene expression data, it is not designed with interventional contexts in mind, but primarily for observational data. This data is less informative for predicting edge directions but more commonly available.

At the center of GENIE3 is the idea to decompose the network inference over all $p$ features (genes) into $p$ separate problems, where each subproblem consists of independently identifying the regulators of a gene.

Fundamentally, one assumes that for each gene $x_j$ there is a function for its regulation which explicitly depends on its direct regulators:

$$x_j = f_j(\{x_{\mathbf{PA}_j}\}) \tag{A1}$$

Here $x_{\mathbf{PA}_j}$ corresponds to the parents of $x_j$. The approach to identifying regulators is to search for genes whose expression values are predictive of that of the target gene - this can suggest a regulatory influence. For each gene potential regulators are weighted by their ability to predict the target gene and ranked from most to least relevant.

For this purpose, the authors chose a tree-ensemble-based method (random forest) for among others its ability to handle a large number of input features, fast computation, being parameter-free and handling interacting and non-linear regulation.

In each subproblem $j$ ($j = 1, ..., p$) Decision Trees are trained on a sample-subset to approximate the regulation function for the gene $X_j$:

$$x_j \approx \tilde{f}_j(\{x_1, ...., x_p\} \setminus \{x_j\}) \tag{A2}$$

Once a tree approximating this function has been computed it is possible to use a *variable importance measure* to quantify the predictive power of each input variable $x_i$ on the expression of gene $j$. These importance measures are then averaged over trees in the ensemble to give a ranking of likely regulators for the gene $j$.

With the likely regulators for each gene $j$ identified by a weight-score in each subproblem, the complete regulatory network can be 'stitched together' by combining all the weighed regulatory edges. In practice some value for a cutoff then has to be chosen, where all edges above this cutoff-value then produce a directed graph representing the GRN.

#### A.2 GRNBoost2

GRNBoost2 [3, 20] is derived from GENIE3, but includes a number of modifications to the tree-ensemble-computation that drastically increase computation speed as well as scalability. As there is little change to the design concept of GENIE3 the quality of inference results is largely the same.

Instead of simple fixed decision trees it employs Gradient Boosting. With this it trains decision trees on randomly sub-setted observations, continually estimating the performance on remaining observations and stopping once improvement drops off. This allows it to avoid a large amount of unnecessary computational load on regressions that have plateaued / ceased improving early on.

#### A.3 Mean Difference

For this approach simply the total causal effect of each intervened gene to others is estimated. To do this expression levels of normalised counts in the unperturbed and perturbed settings are averaged and their absolute difference is computed for each intervention target:

$$|\bar{x}_{i,obs.} - \bar{x}_{i,interv.}| \tag{A3}$$

This value is then used as an edge weight for the edge from the targeted feature to feature $i$.

#### A.4 Greedy Interventional Equivalence Search (GIES)

GIES is a score-based structure learning algorithm that extends a previously existing Greedy Equivalence Search (GES) method to consider interventional contexts as well as observational contexts, developed by Hauser and Bühlmann [12]. It operates on sets of graphs which are not mutually distinguishable by their entailed distributions, referred to as Markov equivalence classes, and which are represented by partially directed graphs containing both directed and undirected edges (essential graphs) [27]. In the space of essential graphs it iteratively searches for a configuration that maximises a score function with respect to the data. For this it fits a linear Gaussian structural causal model to the data, which can be formulated in the following way:

$$X_i = \sum_{j=1}^{d} \beta_{ij} X_j + \epsilon_i, \quad \epsilon_i \sim \mathcal{N}(0, \sigma_i^2), \quad 1 \le i \le d \tag{A4}$$

where $\beta$ is the weighted adjacency matrix, which is constrained to the current essential graph $\mathcal{D}$ with $\beta_{ij} = 0$ if $j \notin \mathrm{pa}_{\mathcal{D}}(i)$, $\sigma_i$ are the feature-specific variances. With this formulation the parameters $\beta$ and $\sigma$ are optimized to maximize the likelihood of the model $\ell_{\mathcal{D}}$, which is also equivalent to a minimization of the residual sum of squares for each feature [13]. And finally with the fitted parameters the score $s$ of the CPDAG with respect to the samples can be evaluated. By default the Bayesian Information Criterion (BIC) is used, which is based on the Log-Likelihood of the model:

$$s(\mathcal{D}; \mathbf{X}) = sup\{\ell_{\mathcal{D}}(\beta, \sigma^2; \mathbf{X}) | \beta \in \beta(\mathcal{D}), \sigma^2 \in \mathbb{R}^p_{>0}\} - \frac{k_{\mathcal{D}}}{2} \log(n) \tag{A5}$$

The first term denotes the maximum likelihood $\ell_{\mathcal{D}}(\beta, \sigma^2; \mathbf{X})$ of the causal model of Equation A4 with respect to the data $\mathbf{X}$. The second term is a penalty term proportional to the number of model parameters $k_{\mathcal{D}} = d + |E_{\mathcal{D}}|$, where $E_{\mathcal{D}}$ is the edge set of $\mathcal{D}$. It punishes adding a large number of parameters to the model, which might result in overfitting.

In order to optimize the CPDAG, GIES performs a greedy search with three different

phases. In the forward phase it explores adding directed edges to improve the score, while maintaining acyclicity. Subsequently in the backward phase it prunes edges which improve the score when removed. And finally in the turning phase it considers all the edges present in the graph, and examines whether reorienting their direction without modifying the skeleton of the graph improves the score.
In each of these phases the evaluation of whether a change improves the score $\Delta s > 0$, can be carried out in parallel for many modifications of the current graph, and if any change is found to be favourable, it is applied and the search continues from this new graph. This is done until in each phase no optimization can be made, at which point the resulting graph is returned as the inferred Causal Graph.

#### A.5 Differentiable Causal Discovery with Interventions (DCDI)

DCDI [4] is a method for causal structure learning that models both observational and interventional data using gradient-based optimization. It extends the earlier differentiable causal discovery algorithm NOTEARS by Zheng et al. [29] to interventional settings and thus improving identifiability. By formulating the structure learning problem as a continuous optimization task, DCDI provides a scalable alternative to traditional combinatorial approaches such as GIES.
In general, Differentiable Causal Discovery algorithms circumvent the expensive combinatorial search over the super-exponential number of DAGs (with respect to the number of variables) by reformulating the optimization problem.
DCDI jointly learns both the structure of the causal graph $G$ and the functional assignments $f_i$ for each variable $x_i$. To enable this, the discrete adjacency matrix representing the graph is relaxed into a continuous form by assigning each edge a weight between zero and one. These weights can be interpreted as edge probabilities or strength of influence.

Rather than iterating over individual graph candidates, DCDI learns the graph structure and conditional distributions simultaneously through gradient-based optimization of a score function. Initially, the model allows interactions between many variables, but applies increasing regularization over time to encourage sparsity and recover a minimal causal graph consistent with the data.
In the following, we first describe how DCDI parametrizes both the causal graph and the conditional distributions of each variable. We then turn to the optimization procedure used to estimate these parameters under structural constraints.

##### Parametrization

In order to learn the causal structure DCDI models both the causal graph and the distributions in a differentiable way.
Each variable $x_i = f_i(x_{pa_i}, N_i)$ is modelled with a normal distribution similarly to GIES in the default configuration. But in favour of more expressiveness the parameters $\mu, \sigma$ are modelled as a non-linear function $f_i$ of the parent features as opposed to a linear one. As a general non-linear function approximator a multi-layer perceptron (MLP) is used:

$$x_i \sim \mathcal{N}(\mu_i(x_{pa_i}), \sigma_i(x_{pa_i})) \tag{A6}$$

$$(\mu_i, \sigma_i) = MLP(x_{pa_i}; \phi_i^{obs.}) \tag{A7}$$

Where $\phi_i^{obs.}$ are the parameters of the MLP which are optimized during training. In order to make this formulation differentiable with respect to the causal graph $G$, the boolean adjacency matrix $A$ is derived from an underlying weighted adjacency matrix $\Lambda$. Specifically, each element of the boolean matrix is modelled as Bernoulli-distributed with the probability rate $\Lambda_{ij}$.

$$A_{ij} \sim Bern(\Lambda_{ij}) \tag{A8}$$

Each individual feature then follows a normal distribution with the following mean and variance::

$$(\mu_i(x), \sigma_i(x)) = MLP(A_i \odot x; \phi_i^{obs.}) \tag{A9}$$

Here $A$ is the binary adjacency matrix, and thus $A_i \odot x$, the Hadamard Product of a column of the matrix with the variable values, masks non-parents of node $i$ and feeds values of selected parents to the neural network. The neural network then outputs the parameters of the density function $(\mu, \sigma)$ for each feature. The density function is a normal distribution $\mathcal{N}(\mu, \sigma^2)$ with parameters mean and standard deviation by default, but it can be extended to any other density function in principle, we explore some other distributions in Section B.

The altered probability density of an intervened on gene $i$ is modelled by replacing the learned conditional distribution $\mathcal{N}(\mu_i(x), \sigma_i(x))$, for this the observational learned NN parameters $\phi_i^{obs.}$ are replaced with interventional parameters $\phi_i^{interv._k}$ specific to each intervention $k$. Thus for each intervention a separate distribution is learned for the targeted genes - while based on the assumption of independent causal mechanisms, all other genes $j$ are still modelled in the original form with $\phi_j^{obs.}$.

Thus for a set of intervened on genes $\mathcal{I}_k$ the full joint probability distribution of DCDI could be written in a factorized form as:

$$\mathbf{P}(\mathbf{x}; A_i, \phi) := \prod_{j=1}^{d} p_{\mathcal{N}} \left( x_j; \mathrm{NN} \left( A_j \odot x; \phi_j^{(k_{j\in\mathcal{I}_k})} \right) \right) \tag{A10}$$

where $p_{\mathcal{N}}$ is the normal probability density and $k_{j\in\mathcal{I}_k}$ is zero if $j$ is not a target of the intervention, and one otherwise.

#### Optimization

The parameters of DCDI, which include the weighted adjacency-matrix elements and the NN-parameters for each features distribution, can be optimized using back-propagation and gradient descent on a score $\mathcal{S}$ which is formulated as the log likelihood

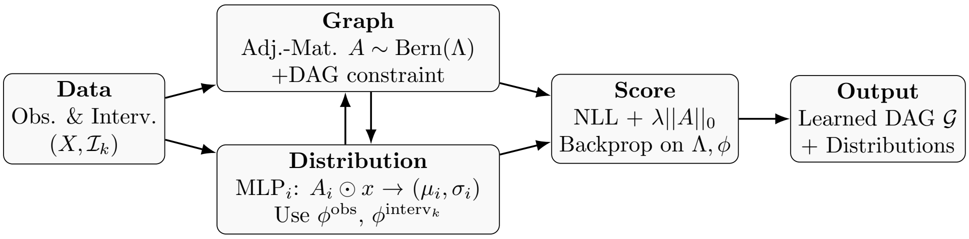

**Fig. A1** Schematic overview of the DCDI model. Observational and interventional data are used to jointly learn the structure of a causal graph (via a parametrized adjacency matrix $A \sim \text{Bern}(\Lambda)$) and the conditional distributions of each variable (via MLPs). These components are optimized jointly via backpropagation on a score that includes log-likelihood, sparsity, and acyclicity penalties.

plus a regularization term:

$$\mathcal{S} := \sum_k \underset{A \sim \text{Bern}(\Lambda)}{\mathbb{E}} \left[ \underset{X \sim p^{(k)}}{\mathbb{E}} \log f^{(k)} \left( X; A, \phi^{(k)} \right) - \lambda ||A|| \right] \tag{A11}$$

Here $f^{(k)}$ are the probability densities and $\lambda$ is the weight of the regularisation penalty on the number of edges in the graph, incentivising the model to recover the simplest graph that explains the data (i.e faithfulness, see [22]). For optimization the expectation value of the final score over each of the outcomes $A_{ij} \in \{0, 1\}$ of the adjacency matrix is maximised. In addition to the score $\mathcal{S}$, a constraint is applied on the adjacency-matrix, to ensure that the resulting graph is acyclic. This is achieved using a continuous constraint on the trace of the matrix exponential, which is satisfied if and only if $A$ represents a DAG [29].

$$\max_{\Lambda, \phi} \quad \mathcal{S} \quad \text{s.t.} \quad \text{Tr} e^A - d = 0 \tag{A12}$$

To optimize $\mathcal{S}$ under this constraint, DCDI uses an augmented Lagrangian method, and gradually introduces both the acyclicity constraint and the sparsity regularization during training to improve stability.
The functional principles of how DCDI is used are also schematically shown in Fig. A1. Finally, it might be worth noting that there is DCD-FG [17], which is a follow-up model that builds upon DCDI and introduces several modifications aimed at improving scalability and practical performance. Most notably, it substitutes the full DAG over $d$ observed variables with a low-rank representation in the form of so-called factor graphs, defined over a smaller set of $m \ll d$ factor vertices, which in turn map to all $d$ variables. In addition it replaces the acyclicity constraint based on the matrix exponential trace with a low-rank variant, such as a spectral radius or trace-exponential criterion applied to a smaller matrix, which is computationally more efficient to evaluate. These structural restrictions significantly reduce the search space and computational cost, enabling the model to scale to datasets with thousands of variables, beyond the typical limits of DCDI, which are on the order of a hundred variables.

#### A.6 Bicycle

Bicycle [24] draws in part on the approach of differentiable causal discovery used by DCDI, but makes different choices in how probability distributions and the adjacency matrix are modelled. In particular it eliminates the requirement of acyclicity of the SCM and instead makes a different assumption - it assumes that the observed gene expression data stems from the steady state distribution of an Ornstein-Uhlenbeck process.

$$dz_i(t) = \left( \alpha_g + \sum_{i \neq h} \beta_{hi} z_h(t) - z_i(t) \right) dt + \sum_h \sigma_{ih} dW_h(t) \tag{A13}$$

Under this assumption the steady state observed data follows a multivariate normal distribution $\mathcal{N}(\overline{z}, \omega)$, for which the parameters mean $\overline{z}$ and covariance $\omega$ directly result from the SDE parameters [26, 28]:

$$\begin{aligned} & z \sim \mathcal{N}(\overline{z}, \omega) \\ & \overline{z} = (B)^{-1}\alpha, \quad \text{where} \quad B = \mathbb{I}_d - \beta^T \\ & \omega \quad \text{s.t.} \quad B\omega + \omega B^T = \sigma\sigma^T \end{aligned}$$

Additionally, Bicycle explicitly models technical noise/measurement error of the observed counts with the multinomial discrete probability distribution, where the mean (probability of each feature) follows the underlying multivariate normal of the SDE.

As such Bicycle is trained by optimizing the parameters of the Ornstein-Uhlenbeck SDE to fit the model's distributions to the measurements. The parameters are given by a set of matrices, and interventions are modelled by replacing the rows of these matrices for the targeted genes with separate parameters.

This introduces an additional layer to the model where the probabilities $p$, i.e. the parameters of the multinomial, are optimized with respect to the observed counts $X$, and the parameters of the SDE are optimized such that their induced multivariate normal $\mathcal{N}(\overline{z}, \omega)$ matches the probabilities of the multinomial.

$$X \sim Mult(n = l_j, \mathbf{p} = \text{Softmax}(z))$$

Where $l_j$ is the library size of cell $j$, which is the total count number the multinomial is conditioned on. Bicycle uses stochastic variational inference to optimize the parameters of the SDE, and maximum likelihood estimation to optimize the probabilities of the multinomial. It employs regularization on the $l_1$-norm of $\beta$ to incentivize sparsity.

#### A.7 Amortized Variational Causal Discovery (AVICI)

AVICI by Lorch et al. [18] is a supervised machine learning approach to causal structure discovery that formulates the task as an amortized inference problem. Instead of performing an often costly search over possible causal models, it learns to predict causal graphs directly from data using a variational inference model, which is trained on synthetically generated data with corresponding ground truth causal graphs.

##### Learning Objective

The amortized inference approach is formulated with the posterior density of causal graphs $p(G|D)$ as a learning objective; for this it is assumed that the data generating distribution $p(G)$, from which samples are available, can be decomposed into an underlying distribution of causal structures $p(G)$ and a data-generating mechanism $p(D|G)$.

To approximate this posterior $p(G|D)$ the forward Kullback-Leibler Divergence $D_{KL}(p|q)$ between an estimate and the true underlying posterior is minimised.

$$\min_{\phi} \quad \mathbb{E}_{p(D)} D_{KL}(p(G|D)||q(G; f_\phi(D))) \tag{A14}$$

Using the definition of the KL Divergence and the Bayes' theorem this objective can also be rewritten in a tractable manner, where the intractable posterior $p(G|D)$ is replaced by a maximisation of the expectation over the available graphs $G \sim p(G)$ and resulting samples $D_G \sim p(D|G)$:

$$\max_{\phi} \quad \mathbb{E}_{p(G)} \mathbb{E}_{p(D|G)} [\log q(G; f_\phi(D))] \tag{A15}$$

Where $q(G; f_\phi(D)) \approx p(G|D)$ is the learned density which approximates the posterior. Using synthetic data a graph is generated using a model $G \sim p(G)$ and then the graph is used to generate samples with a specified data-generating process $D_G \sim p(D|G)$. These pairs $(G, D)$ are then used to compute an empirical expectation value of Equation (A15), which is maximised to optimize the approximation $q(G; f_\phi(D))$.

For scenarios where causal graphs are assumed to be DAGs, AVICI includes a differentiable acyclicity constraint based on the spectral radius of the adjacency matrix in the training objective.

##### Architecture

The inference model $f_\phi$ is a neural network which maps an input dataset $D \in \mathbb{R}^{n \times d}$ ($n$ samples, $d$ variables) to a weighted adjacency matrix $\theta_{d \times d}$, where, similarly to DCDI, each element $\theta_{ij}$ indicates the probability of a directed edge $i \to j$ in the Graph $G$:

$$f_\phi : \mathbb{R}^{n \times d} \to [0, 1]^{d \times d} \tag{A16}$$

$$g_{ij} \sim Bern(\theta_{ij}), \quad \theta = f_\phi(D) \tag{A17}$$

To model $f_\phi$ AVICI uses an architecture designed to respect the invariances of the problem, and which can adapt to datasets of different sample and variable numbers.

To achieve this it maps the input to a tensor of increased dimensionality $n \times d \times k$, and performs multi-head self attention in an alternating fashion. First attention over the variables $d$, which accounts for interactions between the features, followed by attention over the sample axis, to capture variation between samples, including changes of variables under interventions, is taken.

In the final step, the latent dimension $k$ is again collapsed to produce two $n \times d$ embeddings. After normalization the inner product of these two embeddings is then taken to compute the output edge probabilities $\theta_{d \times d}$.

The model architecture is based on a deep, permutation-invariant attention mechanism. The input is mapped to a tensor of shape $n \times d \times k$, where $k$ is the latent feature dimension. Each of the $L$ model layers performs a) self-attention across the variables $d$ to account for interactions between the variables, and b) self-attention across the samples, to capture variation between samples, including changes under interventions.

After the attention layers the model collapses the sample axis $n$, computing two normalized embeddings $\mathbf{u}_i, \mathbf{v}_j \in \mathbb{R}^k$ for each variable, which are used to compute the edge probabilities with an inner product:

$$\theta_{ij} = \sigma\left(\tau\, \mathbf{u}_i^\top \mathbf{v}_j + b\right) \tag{A18}$$

where $\sigma$ is the logistic function, and $b$ and $\tau$ are learned parameters. Since all layers and operations, including the multi-head self-attention and inner product, respect the problem's symmetries, the final output is permutation invariant over samples, and permutation-equivariant over variables.

### B Adaptation of DCDI to Noise

In order to optimize DCDI for the context of scRNA-seq data, we choose to modify its likelihood distribution from the default Gaussian Likelihood to one more appropriate for biological gene expression data. When tackling this, we first have to consider which transformations to apply to the data, and then choose a probability distribution that is able to represent the data distribution under the chosen transformation. Thirdly, the option remains of how to parametrize the distribution, where we should consider to which degree parameters should be free or constrained, and which parametrisation might simplify the optimization during training.

Choosing a few different approaches in each of these options, we constructed a set of modified DCDI models, which we then applied to evaluate their performance.

As data transformation for one we choose to follow common practice of normalizing the sequencing depth of the cells (number of UMI-counts per cell) and to transform the data using a shifted logarithm (as described in section 5.2). This results in a data distribution which includes a fraction of zero-counts, and remaining values distributed in a bell-distribution around a non-zero mean $\mu > 0$. In addition, we also designed a model to be applied to the untransformed discrete UMI Count data.

#### Distribution choice

1. As a probability distribution to model the transformed data we used a mixture of two Gaussian distributions:

$$\mathcal{L} = \prod_i (1 - p_{\mathrm{drop},i}) \times \mathcal{N}(x_i; \mu_i, \sigma_i) + p_{\mathrm{drop},i} \times \mathcal{N}(x_i; 0, 0.1) \tag{B19}$$

One Gaussian with a variable $\sigma$ and $\mu > 0$ is intended to model the bell-distributed non-zero counts, while the second component with a narrow standard deviation $\sigma = 0.1$ and $\mu = 0$ is intended to account for the zero-counts.

2. Negative binomial probability distribution (DCDI-NB)

To model the 'raw' untransformed data we choose a negative binomial (gamma-Poisson) distribution, which can be viewed as a generalization of a Poisson-distribution where a dispersion $\phi \geq 0$ creates a broader shape with variance higher than a ordinary Poisson distribution. This distribution is commonly used in literature to model the discrete UMI counts in scRNA-seq data.

$$\mathcal{L} = \prod_i NB(x_i; s_j \cdot \mu_i, \phi_i) \tag{B20}$$

instead of the $\mu$, $\sigma$ of the Gaussian distribution, DCDI now predicts $\mu$ and $\phi$ (overdispersion) of the negative binomial distribution. Each mean $\mu_i$ that is predicted by DCDI is also multiplied with a cell-specific size factor $s_j = \frac{l_j}{\mathrm{Med}_k(l_k)}$ (see also Section F.1), which corrects for cell-to-cell variation due to changing sequencing depth,

#### Parametrisation

When choosing parametrization for the mixture model (B19) we kept the identical parametrization of $\mu, \sigma$ and tested a number of ways to parametrize $p_{\mathrm{drop}}$.
a) constant over samples $p_{\mathrm{drop}_i}$ (DCDI-Drop-global)
b) variable over samples $p_{\mathrm{drop}_{ij}}$, learned by DCDI as an additional likelihood parameter (DCDI-Drop-local)
Additionally, we implemented a method where dropout-rate is learned consistently over features by leveraging the mean-sparsity dependence of each distribution. As indicated in Fig. B2 there is a sigmoid-like dependence of mean and sparsity in scRNA-seq data as a result of zero-counts increasing with lower expression (DCDI-Drop-sigmoid). Leaning in part on a previous method proDA [1], which makes use of such an effect in mass spectrometry, we regress a sigmoid function to the mean-sparsity distribution. We can then use this mean-sparsity dependence to obtain the mixture parameter $p_{\mathrm{drop}}$ from the mean $\mu$:

$$p_{\mathrm{drop}}(\mu_{i,j}) = 1 - \sigma(-k(\mu_{i,j} - b)) = \frac{1}{1 + e^{-k(-\mu_{i,j}+b)}} \tag{B21}$$

where $k$ and $b$ are the function parameters which are identical for all genes and samples the model is applied on.

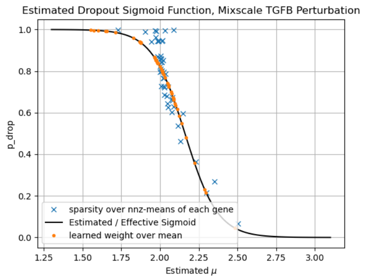

**Fig. B2** Sparsity of each gene over non-zero (nnz) means of each gene observed in shifted-log transformed experimental data (blue, Jiang et al. [16], TGFB Perturbations, A549 cell-line), and mean and mixture rate of Gaussian mixture distributions for each gene that were fitted to the data with the mixture rate given by a sigmoid curve (orange).

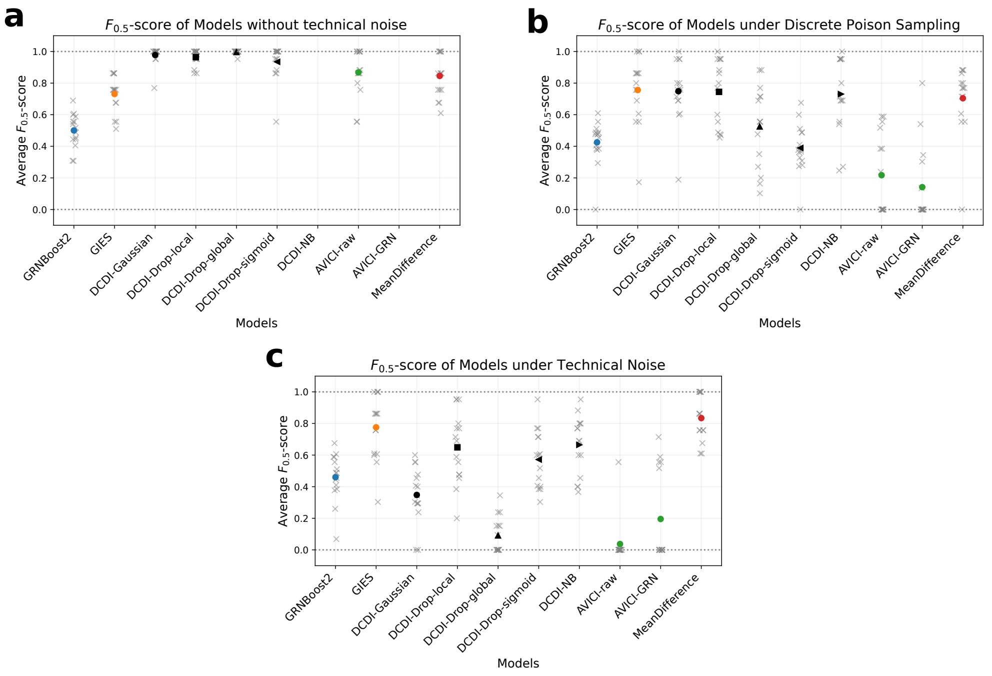

**Fig. B3** $F_{0.5}$ score of all considered DCDI Models when operating on 'raw' data and each of the technical noise transformations. Grey crosses indicate the values produced from each of the 15 datasets, coloured dots indicate the mean over the 15 values. DCDI-NB and AVICI-GRN are only present for two runs, as they are only applicable on discrete count data.

### C Methods: Application of Causal Discovery Algorithms

All methods were run on the same compute cluster with an AMD EPYC 7513 CPU with up to 400GB accessible memory, and access to a GPU for the methods which benefit from it: either an Nvidia A40 (48GB VRAM) or an Nvidia A100 (40GB VRAM). The runtime limit was set at 48h for each GRN inference.

#### GRNBoost2

We used the python implementation of GRNBoost2 through the `arboreto` package `v0.1.16` (available on PyPI)[20]. In its typical application, GRNBoost2 is supplied with a list of TF genes, which a priori restrict regulatory edges in the graph to only ones going out from these. We opted to not restrict GRNBoost2 to these, as this might skew the comparability to other models.

#### GIES

GIES was originally implemented in the R package `pcalg` by the authors [12], but a python implementation by Juan L. Gamella, which is verified to produce identical results, exists [10]. We used this package `gies v0.0.2` from PyPI, using the default BIC score.

#### DCDI

We used the codebase provided by the original authors in the state of 2024 [4]. The hyperparameters we used were: regularisation $\lambda$: $log_{10}(\lambda) \in \{-3, -2, -1.5, -1, -0.5, 0, 0.5, 1\}$, learning rate $lr = 10^{-3}$, 100,000 iterations for synthetic data, 200,000 for real data, 2 hidden layers with 16 units per layer for the MLP. We selected perfect interventions for synthetic data and imperfect interventions for real data, as well as synthetic data with varying effect sizes. Other hyperparameters were kept to their defaults.

We trained DCDI on 80% of the samples and used the remaining 20% to select the lambda value based on the NLL of the trained model. For this, we either simply chose the model with the lowest NLL loss as recommended by the authors, or plotted the NLL of the trained models over the range of lambda values, and selected models with a compromise of high regularisation and low NLL.

#### AVICI

When applying AVICI we used the `avici` v1.0.5 package from PyPI and models that were already pre-trained on synthetic data by the authors, which are available under https://huggingface.co/larslorch/avici.

We used two different models, one, which we refer to as simply "AVICI" or "AVICI-raw", was pre-trained on a mixture of synthetic SCM data with both linear and non-linear regulation models and Gaussian distributions of each feature. The second model we used, which we refer to as "AVICI-GRN", was pre-trained on gene expression data also generated using SERGIO, with technical noise and knockdowns and a variety of feature numbers and graph sizes. Both the SCM and GRN models were trained on data with acyclic underlying causal graphs, for the SCM case the acylicity constraint was enabled, while for the GRN model it was disabled.

For the output of AVICI we then consider whether reported edge probabilities are larger than $p_{ij} > 0.5$, considering only edges that pass this boundary.

#### Bicycle

As Bicycle, which is designed to handle cyclic GRN structures, was only applied on the experimental dataset, which is also the same as the one to which it was applied by the original authors, we used many parameters that were previously found to perform well. Specifically for the loss function we opt to use a $L_1$ regularisation strength $\lambda_{l1} = 1.0$, a weight of the Lyapunov equality of $\xi = 0.1$, and a weight for the Kullback-Leibler divergence between the observed latent gene expressions and steady-states implied by the SDE $\gamma = 1.0$.

We trained the model on 80% of the samples and used the remaining 20% for validation, e.g. to judge if the model converged, training it for 60,000 iterations with a learning rate of $10^{-3}$.

#### Cutoff choice

In order to construct a binary GRN, which is comparable those of other methods, we have to apply a threshold to the weighted interactions scores produced by GRNBoost2, Mean Difference and Bicycle. In the SCENIC workflow Aibar et al. describe a few approaches to thresholding the weights, e.g. by limiting to scores $> 0.001$ or $> 0.005$ [3]. With the knowledge of the GT-edge count $k$, Patrapa et al. similarly compute the precision of only the $k$ highest weighted edges to the GT in their benchmark [23].

We here additionally constructed the following criteria for determining the cutoff $c$, which maximises the product of effect size of interventions on validation samples given by the Wasserstein distance $W_1$ and the edge count $||G_{\geq c}||_0$ in the given GRN $G_{\geq c}$:

$$c = \arg\max_c \quad W_1(G_{\geq c}; X_{val})^\phi \cdot ||G_{\geq c}||_0 \tag{C22}$$

We applied this measure alongside manually chosen fixed thresholds, chosen with reference to the estimated expected edge counts, as e.g. given by CollecTRI [21] for experimental scenarios. However in experimental data we found the measure of Eq. C22 to produce highly fluctuating cutoff-values due to a high range of effect sizes among the perturbations (See Fig F8), and thus proceeded with the following cutoffs: GRNBoost2: $c$ chosen according to Eq. C22 with $\phi = 4$ on synthetic data, $c = 2$ on experimental data. Mean Difference: $c = 0.5$, Bicycle: $c = 0.25$.

### D Methods: Evaluation Metrics

#### Precision & Recall

To calculate precision and recall for a predicted causal graph, all possible edges are counted ($n(n-1)2$) and categorised into TP, FP, TN and FN. TP is a correctly predicted edge, FP is an incorrectly predicted edge, and FN is not predicting an edge where there should be one. Edge direction is considered if not stated otherwise, i.e. predicting an edge between 2 features with the wrong orientation will count as both a FN and a FP. Precision and recall are then computed, for cases where the predicted GRN is empty we set the precision to zero.

#### $F_\beta$-score

The $F_\beta$ is a generalisation of the F-score. The $F_1$-score ($\beta = 1$) is the harmonic mean of precision and recall, thus combining both into one metric. The $F_\beta$-score applies a factor to the precision, thereby weighing its importance:

$$F_\beta = (1+\beta^2) \cdot \frac{\text{Precision} \cdot \text{Recall}}{(\beta^2 \cdot \text{Precision}) + \text{Recall}} \tag{D23}$$

#### Jaccard-Indices between graphs

A directed graph $\mathcal{G} = (V, A)$ is described by the set of nodes $V$ on which it operates, and a set of ordered pairs $A$ of these nodes, which denote the edges present in the graph. If we have two graphs that operate on the same set of nodes $V$, $\mathcal{G} = (V, A_\mathcal{G}), \mathcal{H} = (V, A_\mathcal{H})$, we can define a Jaccard index between these graphs as the Jaccard index between their sets of edges $A_\mathcal{G}, A_\mathcal{H}$:

$$J(\mathcal{G}, \mathcal{H}) = \frac{|A_\mathcal{G} \cap A_\mathcal{H}|}{|A_\mathcal{G} \cup A_\mathcal{H}|} \tag{D24}$$

This index is one if the graphs are identical and zero if their edges are disjoint.

#### Mean Wasserstein Distance $W_p$

Following ideas from the CausalBench benchmark publication [5], we quantify the quality of the predicted edges using intervention samples. For this, we select all genes $B$ where a direct incoming edge $A \to B$ was predicted, and perturbation samples for $A$ are available. For this we quantify the change of the distribution $P_B^{\mathfrak{C}}$ of $B$ when $A$ is unperturbed to $P_B^{\mathfrak{C};doA=X}$ when $A$ is perturbed, using, e.g. the Wasserstein distance.

$$< W_1(\mathcal{H}; X) > := \frac{1}{|\mathcal{I}|} \sum_{i \in \mathcal{I}} \frac{1}{|\mathbf{CH}_i^\mathcal{H}|} \sum_{j \in \mathbf{CH}_i^\mathcal{H}} W_1(\tilde{\mu}_j, \tilde{\mu}_j^{\text{do } X_i}) \tag{D25}$$

#### Precision and Recall of predicted GRNs with respect to Biological Annotation

As biological annotation we opted to use a not cell-type specific curated GRN for Homo Sapiens, CollecTRI, which is compiled from a large number of experimentally verified regulatory interactions, and comes in the form of graph, identifying all TFs and their respective regulated genes [21]. Based on this we can compute a precision and recall of a GRN to the CollecTRI GRN, taking only those biological edges into account that are between the genes being considered in the GRN inference.

#### Comparison against DE-genes

We can compare whether the predicted causal graphs match the pattern of differential expression (DE) visible in the data under interventions.

In order to achieve this, we take each intervention which has samples in the test data and collect all not-targeted features which exhibit differential expression under perturbation of the target. To do this, we use an adjusted t-test on the normalized expression values of unperturbed and perturbed cells, to compute a p-value for differential expression being present under each perturbation.

In order to control the false discovery rate under this large number of tests $m = n_{feat}(n_{feat} - 1)$ we use the Benjamini?Hochberg procedure at a level of the FDR of $\alpha = 0.1$. The gene pairs with p-values rejected under this control are then considered, we collect these pairs of significant DE genes in an adjacency matrix $A_{DE}$ and compare this to the predicted causal graph.

$$A_{DE,ij} = \begin{cases} 1, & \text{if } p_l \leq p_{(k)} \\ 0, & \text{otherwise} \end{cases} \tag{D26}$$

where $p_l$ for $l \in \{1, ..., m\}$ is the p value corresponding to perturbation $i$ and gene $j$. Since this DE test will show direct and mediated effects in the same manner but a causal graph will only contain edges for direct interactions, a simple comparison of the correct causal graph to this matrix would throw up false negatives. To account for this, we expand the predicted causal graph $A$ by adding gene pairs indirectly related over $n_{ind}$ intermediate steps as direct edges to it, which we call $A_{n_{ind}}$.

$$A_{n_{ind}} = \bigcup_{i=1}^{n_{ind}} A^i \tag{D27}$$

This expanded causal graph converges to the transitive closure $A^+$ of $A$, that is the graph containing all edges $(x, y)$ if there was a directed path from $x$ to $y$ in $A$, for large $n_{ind}$. We then compare this $A_{n_{ind}}$ to the observed DE-gene pairs $A_{DE}$.

### E Methods: Benchmark with Synthetic Data

#### Data generation using SERGIO

##### Simulation Framework of Gene Expression based on SERGIO

We simulate gene expression data making use of the SERGIO Single-Cell Expression Simulator framework v1.0.0, which is described in its publication, and has the codebase available on github [7]. Here the gene expression is modelled using the Chemical Langevin Equation [7, 11, 25], where gene production rates depend on the expression levels of their regulators.

##### 1. Graph generation

In order to create simulated gene expression data, we need to a priori specify the graph that the GRN follows. To generate these graphs, we leaned on an Erdős-Rényi model, in which, given a specified number of nodes $d$ and edges $N_{edges}$, the graphs are chosen randomly and uniformly from the set of all graphs with $d$ nodes and $N_{edges}$ edges [8]. To do this for directed acyclic graphs, we randomly picked the edges from the pool of all possible edges while imposing the condition that there are no directed cycles.

##### 2. Modelling with Stochastic Differential Equation

The production rate $P_i(t, \mathbf{x})$, which captures regulatory effects, is in general dependent on the concentration of mRNA species other than $x_i$, which produces the following preliminary SDE:

$$dx_i = (P_i(t; \mathbf{x}) - \lambda x_i)dt + q(\sqrt{P_i(t; \mathbf{x})}dW_t + \sqrt{\lambda_i x_i}dW_t) \tag{E28}$$

where $P_i(t; \mathbf{x})$ is the production rate, $\lambda$ is the decay rate and $q$ is a multiplicative constant that controls the overall stochastic noise amplitude [7, 25].

We then model the production rate $P_i$ of gene $i$ in an additive manner over each of the contributions $p_{ij}$ from the different TF species $j$ that regulate it, which each follow the form of a Hill function $\theta$. The set of genes that regulate gene $i$ is here denoted as $R_i$, which would correspond to its parents $\mathbf{PA}_i$ in the GRN. If gene $i$ has no regulators ($R_i = \emptyset$), $P_i$ can be set to a constant.

$$P_i = \sum_{j \in R_i} p_{ij} = \sum_{j \in R_i} K_{ij}\theta(x_j; h_{ij}, n) \tag{E29}$$

$K_{ij}$ is an interaction weight prefactor that specifies the maximum production rate of gene $i$ induced by each regulating gene $j$.

In application, the interaction weight prefactor $K_{i,j}$ of the SDE was restricted to positive values, which in turn produces only positive regulation contributions $p_{ij}$. From a biological perspective, this means that we model only promoting or up-regulating and not inhibiting or down-regulating interactions. Likewise in the sum of (E29) each term $p_{ij}$ contributes independently and as such only independent TF-target regulatory mechanisms and no combined interactions of multiple regulators on one target are modelled.

#### 3. Sampling Trajectories Over Time

The simulation model so far describes a single cell observed over time. In order to describe the experimental situation where many identical (clonal) cells are measured in the same conditions, SERGIO invokes an ergodic assumption, which postulates that a time average of a system over its trajectory exists, and is the same for almost any initial points. This in turn indicates that sampling over many separate cells which follow the same stochastic equations is equivalent to taking samples from the trajectory of a single cell over time (given that the time intervals are large enough).

In practice, the system is intialised at the mean-reversion level of each gene expression, and then numerically integrated using the Euler-Marayuma method based on Itô Calculus. We let the simulation run for a larger given interval, after which we take a specified number of samples from the trajectory, see Fig. E4 for a characteristic integration trajectory. We repeat this procedure of initialising the simulation and sampling the trajectory a specified number of times, and then all samples are collected in a matrix without regard for their original time ordering.

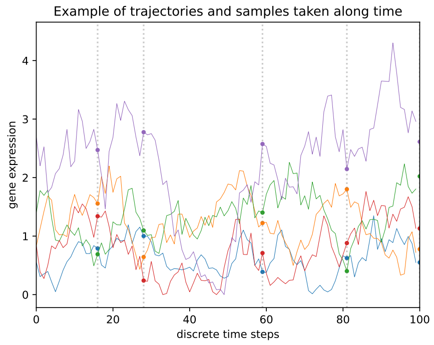

**Fig. E4** Trajectory of 5 genes during a typical Simulation from beginning over 100 discrete time steps. The genes are initialized close to their equilibrium and fluctuate over time, samples are taken at points along the trajectory (dotted vertical lines).

The distribution of the resulting samples appears to closely follow a gamma distribution (Fig. E5). This is also what would be expected from the steady-state distribution of the Cox-Ingersoll-Ross process [6].

#### 4. Simulating Targeted Interventions

With this simulation set-up we then wish to simulate targeted interventions, which can be leveraged for causal learning. For this, an intervention is modelled by modifying the SDE governing the expression of the targeted gene $i$. Specifically, its production rate $P_i(t; \mathbf{x})$ in equation E28 is replaced with a constant term $P_i'$ which can be manually specified by the user or set to a portion of its mean production rate in the unperturbed condition. In practice, we set it to a low percentage of the production rate or to zero.

#### 5. Simulating additional Technical Noise

In addition to simulating the raw, continuous expression data, SERGIO also offers transformations which can be applied to the data to model technical noise that is very prominent in scRNA-Seq data as a result of the measurement process.

The Poisson distribution (under the limit of rare events) is applied to model the observed UMI counts, each simulated continuous concentration is converted to a discrete UMI-count by taking a single sample from a Poisson distribution with that mean.

The dropouts are modelled by multiplying each measured expression with a binary pre-factor sampled from a Bernoulli distribution, setting it to zero with a probability $p_{\text{drop}}(x)$.

$$p(x'|x) = (1 - p_{\text{drop}}(x)) \cdot \text{Pois}_{\lambda=x}(x') + p_{\text{drop}}(x) \cdot \delta(x', 0) \quad \text{(E30)}$$

Where $x$ are the original simulated values without technical noise and $x'$ are the resulting values with technical noise. $\text{Pois}_{\lambda=x}$ is the Poisson distribution with mean $x$.

The Bernoulli dropout probability is distributed following a sigmoid / logistic function of the (logarithmized) expression of the given gene in that cell.

$$p_{\text{drop}}(x_{ij}) = Ber(\pi(x_{ij})) \quad \text{(E31)}$$

$$\pi(x_{ij}) = sig(-k(x_{ij} - x_0)) \quad \text{(E32)}$$

This essentially 'masks out' lower expression values, while leaving others unaltered. The midpoint $x_0$ and $k$ are kept constant over all samples in a dataset, $x_0$ is set to a user-given percentile of expression samples in the data.

The resulting data distributions are characterised in Fig. E5:

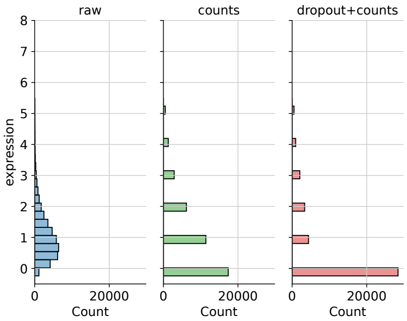

**Fig. E5** Characteristic distribution of data under simulated technical noise

#### Baseline parameter choices

To benchmark with synthetic data, datasets were created using SERGIO ranging from scenarios with simplistic data on which algorithms should perform well to more challenging scenarios which are closer to those in real biological data on which algorithms should be ultimately applied. This should allow to rank each algorithm by how well they perform overall, but also to get a closer view at which features in the data algorithms rely on to perform well and which features might cause some of the algorithms to struggle.

For this purpose, we selected baseline parameters for the data generation, which are outlined in Table E1, which were in part re-used from the SERGIO implementation, and in part selected with consideration to the computational methods.

| **Parameter** | **Value** |
|---|---|
| n feat ($d$) | 10 |
| n interv | 10 |
| Edge Density | 0.5 |
| Samples per run $n_{samp}$ | 200 |
| Number of runs | 20 obs., 5 per interv. |
| dt | 0.1 |
| steps / sample | 15 |
| $q$ Noise Amp.* | 0.6 |
| $\lambda$ Decay Rate | 0.8 |
| Base prod. rate $P_i$* | $[1, 2]$ |
| KD prod. rate $P_i^{KD}$* | $[0.1, 0.2]$ |
| Hill Coeff $n$ | 2 |
| Half response $h$ | mean-exp (unpert.) |
| Interaction strength $K$ | 4 |
| Noise | no |
| dropout_k | 10 |
| dropout percentile | 75 |

**Table E1** Default parameters used in generation of synthetic data. Production rates were sampled uniformly from the given intervals. Stars (*) mark values that were still tweaked during the data generation, and as such differ for some runs.

For each simulation run, the expression values were initiated at the features mean reversion level defined by to Eq. E28, the forward integration was carried out for $n_{samp} \times 15$ time steps and $n_{samp}$ expression values were taken as samples. We first simulated the trajectories of genes at the source of the DAG, and then set the half-response values $h$ of the Hill functions to their respective mean expression when unperturbed.

We ran 20 such integrations in the observational setting, and then five more per perturbed target gene, computing a total of 14,000 samples in the standard setting.

#### Varying parameters - without technical noise

This approach was then applied while varying some of the parameters to create data from a variety of scenarios. Here we did not apply technical noise to begin with, and varied the following data-generation parameters: total sample number, feature number, stochastic noise amplitude, ground-truth graph edge number, intervention number and effect size (knockdown strength). For each of these scenarios, the chosen values are listed in Table E2.

| Run Parameter | Symbol | Values |
|---|---|---|
| Number Samples | $n_{\text{samp}}$ | 20, 100, 200, 400, 800, 3200 |
| Number Features | $d$ | 10, 20, 30, 40, 50, 60, 70, 80, 90, 100 |
| Stochastic Noise | $q$ | 0.1, 0.2, 0.4, 0.6, 0.8, 1.0 |
| Number of Edges | $n_{\text{edges}}$ | 1, 2, 5, 10, 20, 45 |
| Number of Interventions | $n_{\text{interv}}$ | 0, 1, 3, 5, 8, 10 |
| Effect Size | $r_{KD}$ | 1.0, 0.7, 0.4, 0.2, 0.1, 0.0 |

**Table E2** Summarized parameter values used in each of the scenarios

The remaining simulation parameters were kept mostly the same between all these generated datasets. We used a set of 10 features, randomly generated DAGs on these with 5 edges as described previously, and performed interventions on all 10 features.

However, for the following runs, three of the parameters differed from those in Table E1: In the number of edges run the stochastic noise amplitude was set at $q = 0.4$ (instead of $q = 0.6$), in the number of interventions and effect size runs the base production rate was sampled from $P_i \sim \mathcal{U}(0.5, 1.2)$ and in the number of genes run the KD production rate $P_i^{KD}$ was set at 0.1 of the respective mean unperturbed gene expression.

The variation of the stochastic noise amplitude $q$ and effect size $r_{KD}$ produced lower overall effects than $n_{\text{samp}}$, feature number $d$ or $n_{\text{interv}}$, their results are displayed in Fig. E6. The results over varying edge number $n_{\text{edges}}$, with predicted edges split into categories, are displayed in Fig. E7.

#### Varying parameters - with technical noise

In order to benchmark the abilty to handle data with technical noise, we generated data similarly as before under 'easy' conditions for the algorithms (Table E1), and then applied technical noise as described by Eq. E30 to this data. In order generate realistic data with respect to gene mean expression and sparsity we, however, tweaked some parameters from those listed in the table: base production $P_i \in [0.3, 1.3]$, interaction strength $K = 1.8$. We used data with perfect knockouts $r_{kd} = 0.0$ and generated 15 datasets from 15 different randomly sampled DAGs to obtain a larger sample size.

Evaluation was performed on the 15 separate gene expression datasets, where identical data was used first without technical noise and then with **a)** discretised UMI count

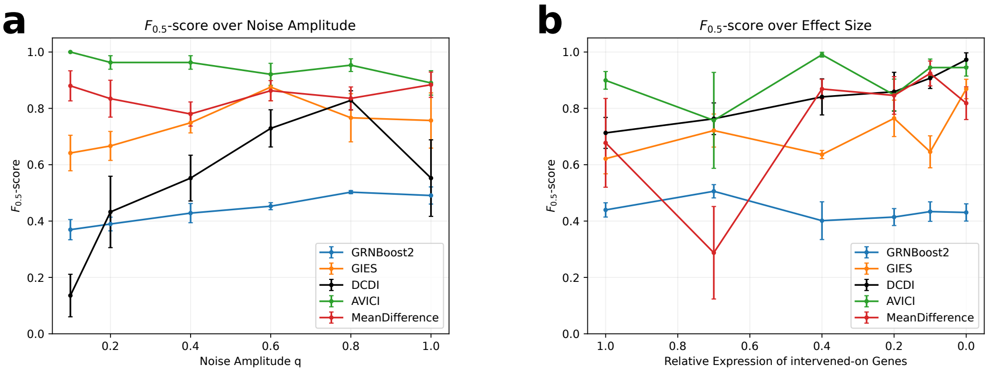

**Fig. E6** Additional results of applying the considered Algorithms on synthetic data in three different Scenarios: a) varying amplitude of stochastic noise and b) varying effects size of interventions

data and **b)** both discretised data and added dropouts, with resulting distributions as characterised in Fig. E5.

### F Methods: Benchmark with Experimental Data

For experimental application, we selected a Perturb-Seq dataset of human-derived melanoma (cancer) cells in which 249 different genes related to immunity over cells in three different conditions were knocked out using CRISPR-Cas9 perturbations [9].

We selected one of the three conditions to obtain a homogeneous pool of cells: Co-culture cells that were cultured with T-cells derived from a patient tumour (73,114 cells). Then, in order to have a dataset of a scale that each of the considered methods can handle, and a high overall quality, we selected 40 genes of the 249 perturbed genes based on the highest number of perturbed cells available in the data for each. This gives us a dataset comprising 40 genes and a total of 12,064 samples, with 3291 unperturbed control samples, and ranging from 137 to 435 samples per perturbation target. The chosen genes with individual cell counts are listed in detail in Table G3.

#### F.1 Data Preprocessing

When working with scRNAseq data, measurements are technical effects include fluctuation of sequencing depth between cells, and a high measurement or 'shot' noise on each value [14]. In order to nevertheless work with this data, a number steps can be taken to alleviate some of the measurement effects [2, 19]. The particular steps we took when preprocessing experimental data were:

**A. quality control**

We removed features which are observed in less than 100 cells, and removed cells that have nonzero UMI-counts observed for less than 500 genes.

**B. transformation of count values**

Then we apply transformations to the remaining UMI counts to correct for measurement effects. First, we normalise the total counts of each cell $l_j = \sum_i x_{ij}$ over the genes $i$ of cell $j$ to those of the median cell $\mathrm{Med}_k(l_k)$ to account for the variation in

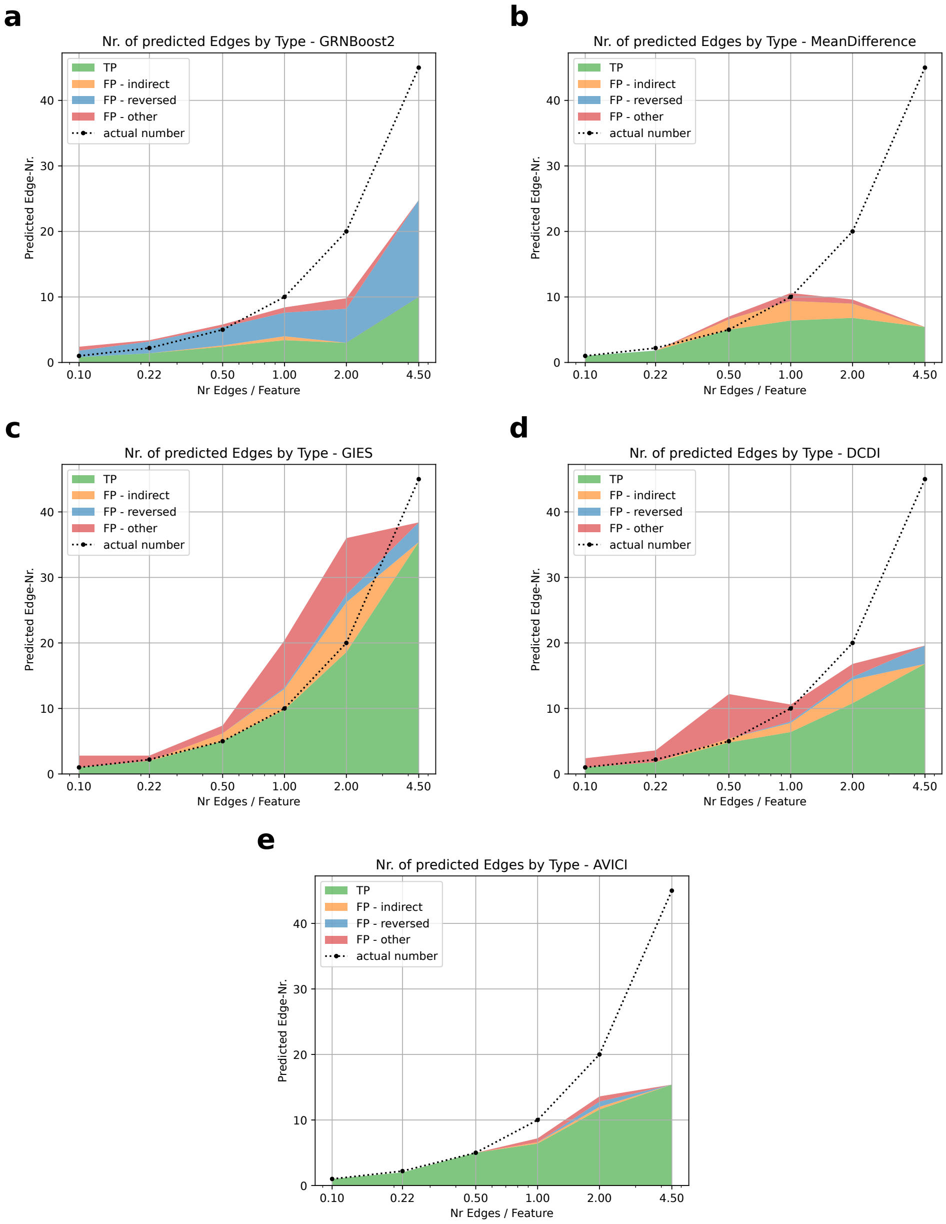

**Fig. E7** Predicted edges categorised by types over data with varying GT edge number: 1) True Positive (TP) (green), correct prediction 2) False Positive (FP) - indirect (orange), predicted edge where there is instead an indirect effect over one in-between node 3) FP-reversed (blue), predicted edge where it would be correct if reversed 4) FP-other (red), false prediction. Values were computed over 5 separate datasets each, plotted are the means for GRNBoost2, Mean Difference, GIES, DCDI and AVICI.

sequencing depth, and then apply a logarithm with an added pseudo-count of 1 to the data. This is intended to counteract the heteroscedasticity of features of different mean expression having significantly different variances, transforming them to have roughly similar variance [2]. The pseudocount accounts for zero-counts, ensuring that they remain at zero. The transformed count value of gene $i$ in cell $j$ is then:

$$x_{ij,\text{transf.}} = \log\left(\text{Med}_k\left(l_k\right)\frac{x_{ij}}{l_j} + 1\right) \tag{F33}$$

To visualise the resulting perturbation data the z-scores of each gene under each perturbation are displayed in Fig. F8.

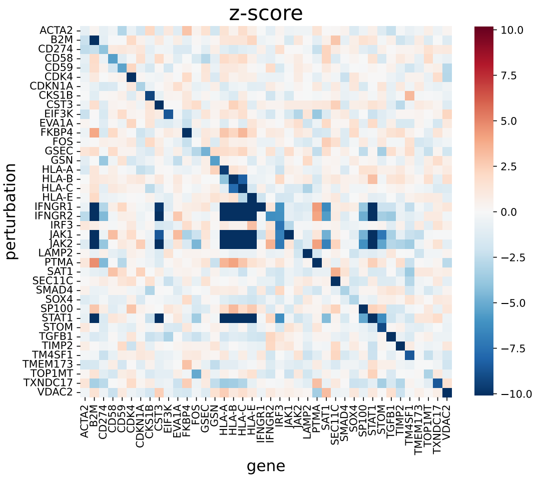

**Fig. F8** Z-scores of each genes expression level under each perturbation for the subset of genes in the experimental data considered. Y-axis indicated the perturbation targets, and x-axis the differentially expressed genes. The diagonal shows down-regulation of target genes, and a number of off-diagonal genes show a very strong reaction.

#### F.2 Application of Causal Discovery Algorithms

In most cases the methods were applied analogously on experimental data as on synthetic gene expression data. The exceptions being that for DCDI-NB we now include the size factor as a model input, which remained set to 1 on synthetic data. For AVICI we now use the GRN model described in section C. Here we further applied Bicycle, which, being designed for unprocessed count gene expression values and cyclic GRNs, is expected to show its strengths primarily here.

### G Additional Results

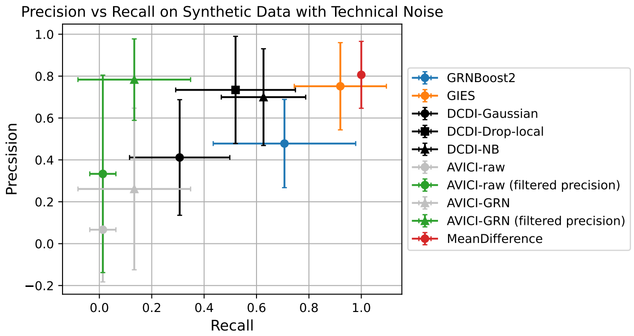

**Fig. G9** Mean precision and recall of the considered methods on synthetic data with technical noise. In grey precision values for empty GRNs (which we set to zero) were included in the mean, in colour precision values of empty GRNs were not considered. Error bars indicate the standard deviation of the mean. AVICI displays a competitive precision, but has a large portion of empty predicted GRNs.

**Table G3** Selected genes and the number of perturbed cells after filtering available for each gene in the used scCRISPR-screen dataset.

| Gene / Perturbation Target | Sample Count |
|---|---|
| non-targeting control | 3291 |
| IFNGR2 | 435 |
| JAK2 | 401 |
| JAK1 | 360 |
| STAT1 | 360 |
| IFNGR1 | 287 |
| B2M | 268 |
| CD58 | 267 |
| CDKN1A | 251 |
| HLA-A | 249 |
| TGFB1 | 247 |
| GSN | 245 |
| ACTA2 | 243 |
| CD59 | 231 |
| LAMP2 | 230 |
| VDAC2 | 229 |
| STOM | 217 |
| PTMA | 216 |
| SP100 | 216 |
| CD274 | 215 |
| IRF3 | 213 |
| TMEM173 | 211 |
| TM4SF1 | 211 |
| SOX4 | 199 |
| SEC11C | 197 |
| EVA1A | 192 |
| TOP1MT | 183 |
| CKS1B | 182 |
| SAT1 | 179 |
| SMAD4 | 178 |
| HLA-E | 178 |
| GSEC | 163 |
| FOS | 156 |
| HLA-C | 156 |
| CST3 | 155 |
| TIMP2 | 147 |
| CDK4 | 147 |
| EIF3K | 141 |
| TXNDC17 | 141 |
| FKBP4 | 140 |
| HLA-B | 137 |

[6] John C. Cox, Jonathan E. Ingersoll, and Stephen A. Ross. A theory of the term structure of interest rates, econometrica 53, 385-407. *Econometrica*, 53(2): 385–407, 1985. doi: 10.2307/1911242.

[7] Payam Dibaeinia and Saurabh Sinha. Sergio: A single-cell expression simulator guided by gene regulatory networks. *Cell Systems*, 11(3):252–271.e11, 2020. ISSN

2405-4712. doi: https://doi.org/10.1016/j.cels.2020.08.003. URL https://www.sciencedirect.com/science/article/pii/S2405471220302878.

[8] Paul L. Erdos and Alfréd Rényi. On the evolution of random graphs. *Transactions of the American Mathematical Society*, 286:257–257, 1984. doi: 10.1090/S0002-9947-1984-0756039-5.

[9] Chris J. Frangieh, Johannes C. Melms, Pratiksha I. Thakore, Kathryn Geiger-Schuller, Patricia Ho, Adrienne M. Luoma, Brian Cleary, Livnat Jerby-Arnon, Shruti Malu, Michael S. Cuoco, Maryann Zhao, Casey R. Ager, Meri Rogava, Lila Hovey, Asaf Rotem, Chantale Bernatchez, Kai W. Wucherpfennig, Bruce E. Johnson, Orit Rozenblatt-Rosen, Dirk Schadendorf, Aviv Regev, and Benjamin Izar. Multimodal pooled perturb-cite-seq screens in patient models define mechanisms of cancer immune evasion. *Nature Genetics*, 53:332 – 341, 2021. doi: 10.1038/s41588-021-00779-1. Single Cell Portal https://singlecell.broadinstitute.org/single_cell/study/SCP1064/.

[16] Longda Jiang, Carol Dalgarno, Efthymia Papalexi, Isabella Mascio, Hans-Hermann Wessels, Huiyoung Yun, Nika Iremadze, Gila Lithwick-Yanai, Doron Lipson, and Rahul Satija. Systematic reconstruction of molecular pathway signatures using scalable single-cell perturbation screens. *Nature Cell Biology*, pages

1–13, 2025.

[22] Jonas Peter, Dominik Janzing, and Bernhard Schölkopf. *Elements of Causal Inference - Foundations and Learning Algorithms*. The MIT Press, Cambridge, Massachusetts, 2017.

[23] Aditya Pratapa, Amogh Prabhav Jalihal, Jeffrey N. Law, Aditya Bharadwaj, and T. M. Murali. Benchmarking algorithms for gene regulatory network inference from single-cell transcriptomic data. *Nature methods*, 17:147 – 154, 2019. doi: 10.1038/s41592-019-0690-6.

[24] Martin Rohbeck, Brian Clarke, Katharina Mikulik, Alexandra Pettet, Oliver Stegle, and Kai Ueltzhöffer. Bicycle: Intervention-based causal discovery with cycles. In Francesco Locatello and Vanessa Didelez, editors, *Proceedings of the Third Conference on Causal Learning and Reasoning*, volume 236 of *Proceedings of Machine Learning Research*, pages 209–242. PMLR, 01–03 Apr 2024. URL

https://proceedings.mlr.press/v236/rohbeck24a.html.
